# Self-Sufficient Maturation and Catalysis of a Clade E CODH Encoded in a CooCTJ-Operon from *Clostridium pasteurianum* BC1

**DOI:** 10.64898/2026.03.10.710785

**Authors:** Maximilian Böhm, Henrik Land

## Abstract

Carbon monoxide dehydrogenases (CODHs) are metalloenzymes central to microbial CO metabolism and CO_2_ fixation. We report the heterologous production and characterisation of *Clostridium pasteurianum* BC1 CODH-III (*Cp*^BC1^CODH-III), from the phylogenetic clade E, co-expressed with its maturation machinery CooCTJ. *Cp*^BC1^CODH-III shows moderate CO oxidation (150 U/mg) and CO_2_ reduction (0.568 U/mg) activities. Electron paramagnetic resonance (EPR) spectroscopy under varying redox conditions identified a rhombic signal at g ≈ 2.0, characteristic of reduced B-clusters, and a C-clusters at different stages (g ≈ 1.75, g ≈ 1.72), indicative of a bound CO_2_. Investigation of maturation effects showed that co-expression of CooCTJ stabilised *Cp*^BC1^CODH-III production, but did not enhance maximum activity, which was primarily influenced by nickel availability. Comparative operon analysis with the well-studied clade F *Rhodospirillum rubrum* CODH (*Rr*CODH) revealed high structural similarity in CODH and CooC, but significant divergence in CooJ, with conserved metal-binding regions identified via AlphaFold3 modelling and dot plot analysis. *Cp*^BC1^CODH-III represents a unique example of a clade E CODH within a clade F genomic context, demonstrating intrinsic robustness in maturation and activity

## Introduction

Nickel-iron carbon monoxide dehydrogenases (CODHs) catalyse the interconversion of CO and CO_2_ at remarkable rates.^[1,2]^ They are central to microbial carbon metabolism and participate in a wide variety of physiological processes, from energy conservation to redox balancing and carbon fixation.^[1,3]^ Despite their overall sequence similarity and structural conservation, CODHs also show diversity in catalytic capabilities. CODHs are classified into several phylogenetic clades (A to G) that generally correlate with distinct metabolic contexts and genomic organisations.^[3,4]^

The overall structure of CODH is homodimeric, with five metal clusters. Three of those are regular [4Fe4S] clusters, called the B-, B’-and D-clusters, while two clusters are biologically unique [Ni3Fe4S] clusters, called C-and C’-clusters. Within the proximity of the C cluster a distal iron is coordinated. Between the nickel and iron, the substrate interconversion is catalysed. Electron paramagnetic resonance (EPR) spectroscopy is an indispensable tool in the study of CODH as it can detect distinct redox states of the metal clusters as well as key intermediates of the catalytic cycle.^[1]^ For example, Basak *et al*. could assign the long known and characteristic signal C_red1_, to a HO-coordinated reduced C-cluster.^[5]^

Due to this complex active site cofactor, CODH-operons often contain putative maturases such as CooC, CooT and CooJ.^[6–8]^ There are however many cases where CODHs are not encoded together with maturases with the most well-studied example being CODH-II from *Carboxydothermus hydrogenoformans* that can reach full activity after expression without any maturases present.^[9]^ Even though the exact mechanism of the maturation process is still unknown, a clear trend is seen in the occurrence of these maturases in CODH encoding operons. In a previous study we showed that while CooC and CooT commonly appear in clade A, E and F, CooJ seems to be almost exclusive for clade E and F CODHs, each with their own operonic arrangements.^[4]^ The occurrence of any maturases in clades B, C, or D is rather scarce, and usually only happens in deeply branching areas of these clades.^[4]^

Nevertheless, our analysis revealed a clade E CODH with a typical clade F-operon arrangement which caught our attention. It is coded within the genome of *Clostridium pasteurianum* BC1, which encodes four CODHs. Two clade C, one clade E and one clade F (Figure 1). This is a relatively high number of CODHs as most organisms only encode one or two.^[4]^ A well-known exception to this is *C. hydrogenoformans* which encodes five CODHs.^[10]^ We suggest the naming *Cp*^BC1^CODH-I (Uniprot ID: R4K676, clade C), *Cp*^BC1^CODH-II (R4K9W7, clade F), *Cp*^BC1^CODH-III (R4K1Q9, clade E) and *Cp*^BC1^CODH-IV (R4KEL0, clade C), with regards to the origin strain and their genome position. The closely related *C. pasteurianum* DMS525 strain only contains two CODH genes, one from clade C (A0A0H3J3W5) and one from clade E (A0A0H3J660).

**Figure 1.**
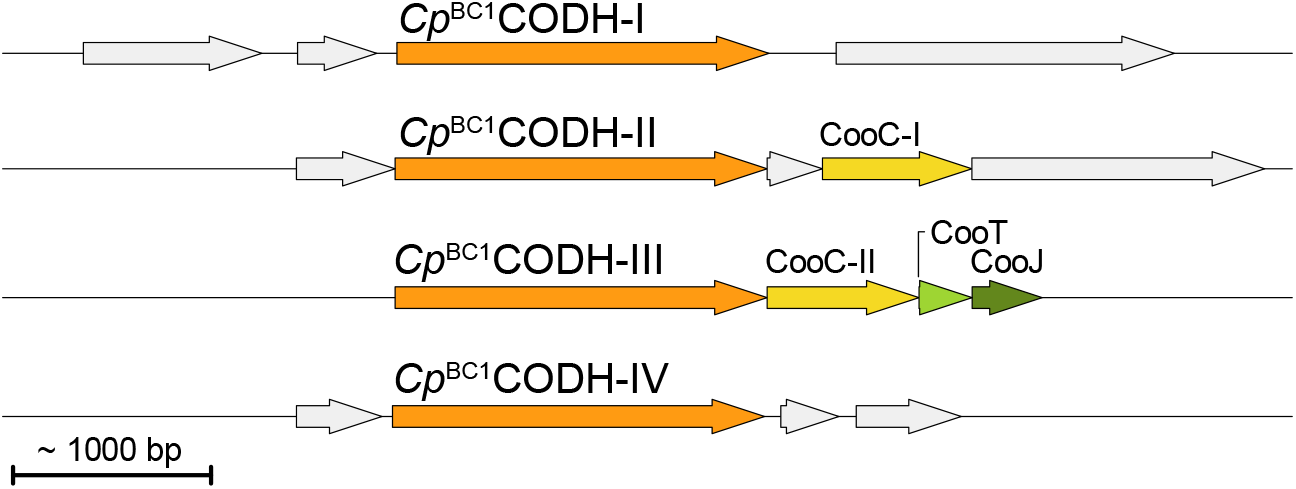
CODH operons of *C. pasteurianum* BC1. From top to bottom from clade C, F, E, and C.

With this in mind, we wanted to investigate how this operon composition, assumed to be clade F specific, affects clade E *Cp*^BC1^CODH-III’s production and activity. For this we produced *Cp*^BC1^CODH-III heterologously in *Escherichia coli* with and without the presence of a maturation machinery and assayed its activity in crude cell lysate. We furthermore purified *Cp*^BC1^CODH-III via Strep-tag II affinity chromatography to determine specific activities and investigate redox states via EPR spectroscopy.

In this study, we show that *Cp*^BC1^CODH-III is catalytically active even when produced in the absence of its associated maturases, indicating intrinsic assembly competence, more affected by expression level and nickel availability. Furthermore, EPR spectroscopy of purified enzyme reveals features consistent with a canonical C-cluster, confirming proper cofactor assembly.

## Results and Discussion

*Cp*^BC1^CODH-III was heterologously produced in *E. coli* with co-expression of its putative maturation machinery CooCTJ. The purified enzymes showed a metal content of 4.5 iron/monomer and 0.7 nickel/monomer. This initial metal content suggested incomplete incorporation of the FeS-clusters, which is common in heterologous expression systems due to differences in the cellular environment compared to the native host. After *in vitro* reconstitution of the FeS-clusters the iron content could be increased to 10 iron/monomer, as expected for a canonical CODH. This result confirmed the successful reconstitution of the FeS-clusters, which are essential for the enzyme’s catalytic activity. The nickel content of 0.7 nickel/monomer is in good agreement with the expected 1 nickel/monomer for a canonical CODH. This near-stoichiometric nickel incorporation indicates efficient nickel insertion, likely facilitated by the co-expressed maturases.

SDS-PAGE analysis showed a band at approximately 70 kDa, corresponding well with the theoretical mass of a *Cp*^BC1^CODH-III monomer with a Strep-tag II (69.9 kDa). Notably, maturation enzymes could not be identified on the gel. This could be attributed to their smaller size or instability under the experimental conditions. The enzyme performed well in CO oxidation and CO_2_ reduction activity assays (Table 1). CO oxidation activity could be increased 19-fold after reconstitution, while CO_2_ reduction activity only increased 1.5-fold. This disparity suggests that the reconstitution process primarily enhances the enzyme’s ability to oxidise CO, possibly due to the restoration of D and B clusters. The modest increase in CO_2_ reduction activity may indicate that efficient electron transfer is not as rate-limiting as it is for CO oxidation. This could be due to low substrate availability, as CO_2_ concentrations in solution might not reach saturation conditions, or a low thermodynamical driving force due the use of methyl viologen as an electron mediator.

**Table 1.** Activity of Cp^BC1^CODH-III towards CO oxidation and CO_2_ reduction. Activity is reported in U per mg of enzyme. One U is defined as conversion of 1 µmol of substrate per minute.

| $Cp^{BC1}CODH$-III | Specific activity [U/mg] | |
| --- | --- | --- |
|  | CO oxidation | CO<sub>2</sub> reduction |
| As prepared | 7.735 ± 0.78 | 0.398 ± 0.009 |
| Reconstituted | 149.7 ± 3.2 | 0.568 ± 0.013 |

When investigating the electronic structure of the *Cp*^BC1^CODH-III metal clusters via electron paramagnetic resonance (EPR) spectroscopy under different redox conditions, we found that all samples show a strong rhombic signal centring around g ≈ 2.0, reminiscent of reduced B-clusters in CODH (Figure 2A). Furthermore, we could observe that these signals remain unaltered when the enzyme is incubated with CO_2_ for 30 min, suggesting a sluggish reaction rate that needs a certain drive to push CO_2_ reduction as seen in solution assays. After incubation with sodium dithionite the rhombic signal increases, likely due to reduction of more enzyme that becomes EPR active. Finally, when incubated with CO, a characteristic change in line shape can be observed when power is increased with two features appearing at low g ≈ 1.75 and g ≈ 1.72 (Figure 2B). This spectral change is characteristic of two C-cluster states: the HO bound state of the reduced C-cluster (C_red1_) and the CO_2_ bound state of the C cluster (C_red2_).^[5]^ Similar mixed features were observed in other CODHs, for example in the CODH from *Nitratidesulfovibrio vulgaris* (*Nv*CODH, formerly known as *Dv*CODH), another clade E CODH.^[11]^

**Figure 2.**
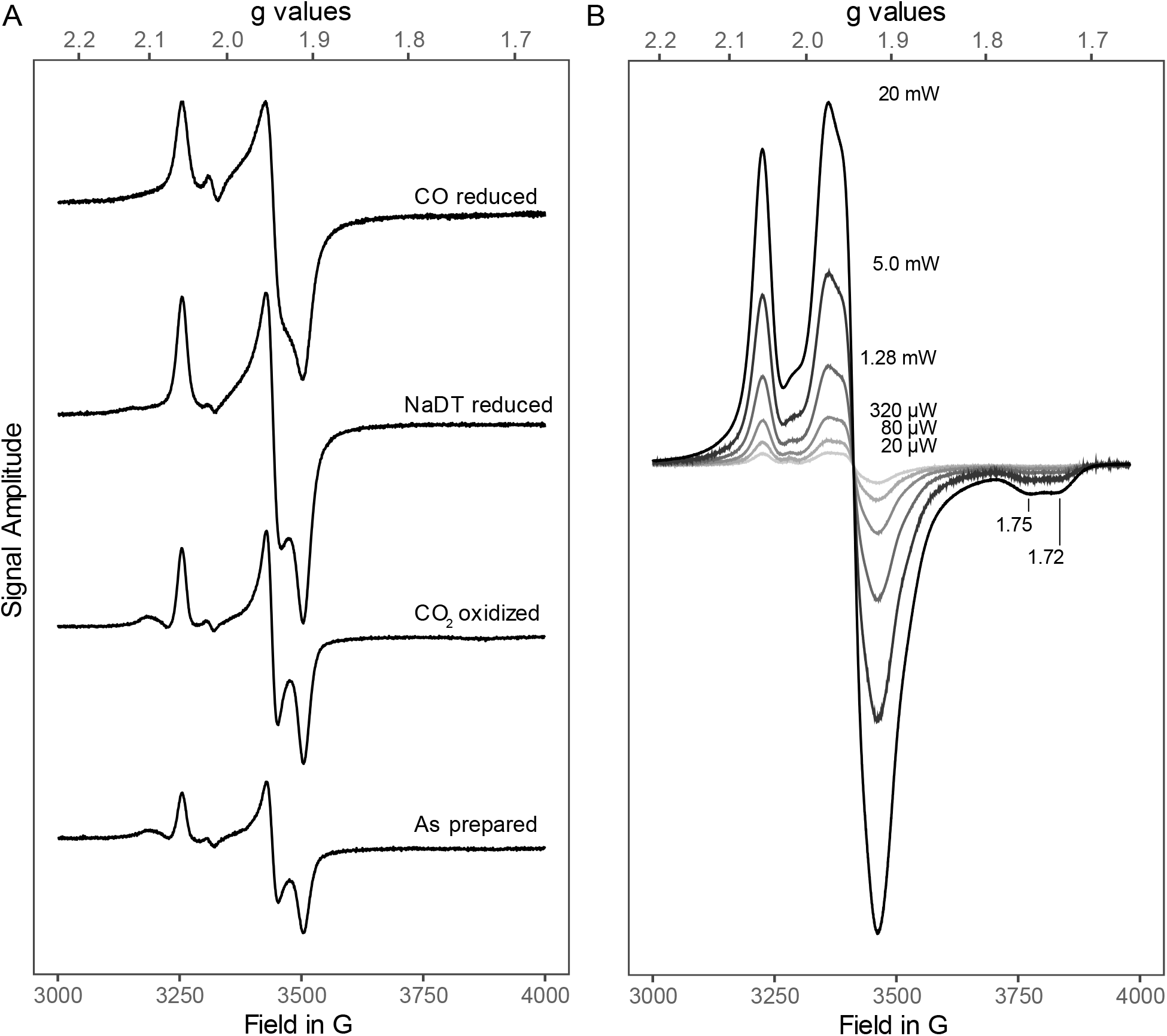
EPR spectroscopy of Cp^BC1^CODH-III in different redox states. A) The protein was incubated with the indicated substrates for 30 min, and then measured at 5 K, 9.37 GHz, 20 µW. B) CO reduced sample was measured at 5 K, 9.27 GHz, at different powers to make the signal in g ≈ 1.75 region more visible.

To understand how maturases influence the production of *Cp*^BC1^CODH-III, we tested the activity of crude cell lysates for CO_2_ reduction after the induction of *Cp*^BC1^CODH-III under different conditions. This approach allowed us to assess the role of maturases in a diverse set of environments. We transformed cells with only CODH or both CODH and maturase plasmids, and selected six colonies each. As a negative control we used *E. coli* with no additional plasmid, to confirm that any observed activity was specifically due to the expression of *Cp*^BC1^CODH-III and not background activity from the host. We tested expression in a 96-deep-well plate with different concentrations of inducer (IPTG) and different concentrations of NiCl_2_ in the medium. After induction the cells were lysed, and the crude cell lysate was used in a 96-well plate CO_2_ reduction assay. The results are summarised in Figure 3. We can see that the maturases have a stabilising effect on the production of active *Cp*^BC1^CODH-III, as IPTG concentration does not significantly affect measured activity when maturases are co-expressed with *Cp*^BC1^CODH-III. This suggests that the maturases help maintain a consistent level of active enzyme, regardless of induction strength, likely by ensuring proper folding and metal incorporation. The overall activity seems unaffected by the maturases, and nickel supply in the medium seems to have a greater impact on the activity of this enzyme, when produced in *E. coli*. This is in stark contrast to other studies on *Rr*CODH that show a clear dependence on both the co-expression of maturases and elevated nickel concentrations for reaching maximum activity, suggesting that *Cp*^BC1^CODH-III is a more flexible enzyme that does not require its maturases.^[12]^ The difference may reflect inherent properties of *Cp*^BC1^CODH-III, such as a higher affinity for nickel or a more robust folding pathway that is less dependent on maturases. Interestingly, other clade E CODHs such as *Nv*CODH, have also been shown to require their maturases to be produced actively.^[11]^ Our findings indicate that the maturases primarily function to stabilise nickel homeostasis in the vicinity of the CODH, minimising fluctuations and thus ensuring more consistent activation across diverse conditions. This could be due to a late acquisition of the maturation machinery, which improved the expression of *Cp*^BC1^CODH-III in its native host, though it was not necessary for an ancestor of this CODH.

**Figure 3.**
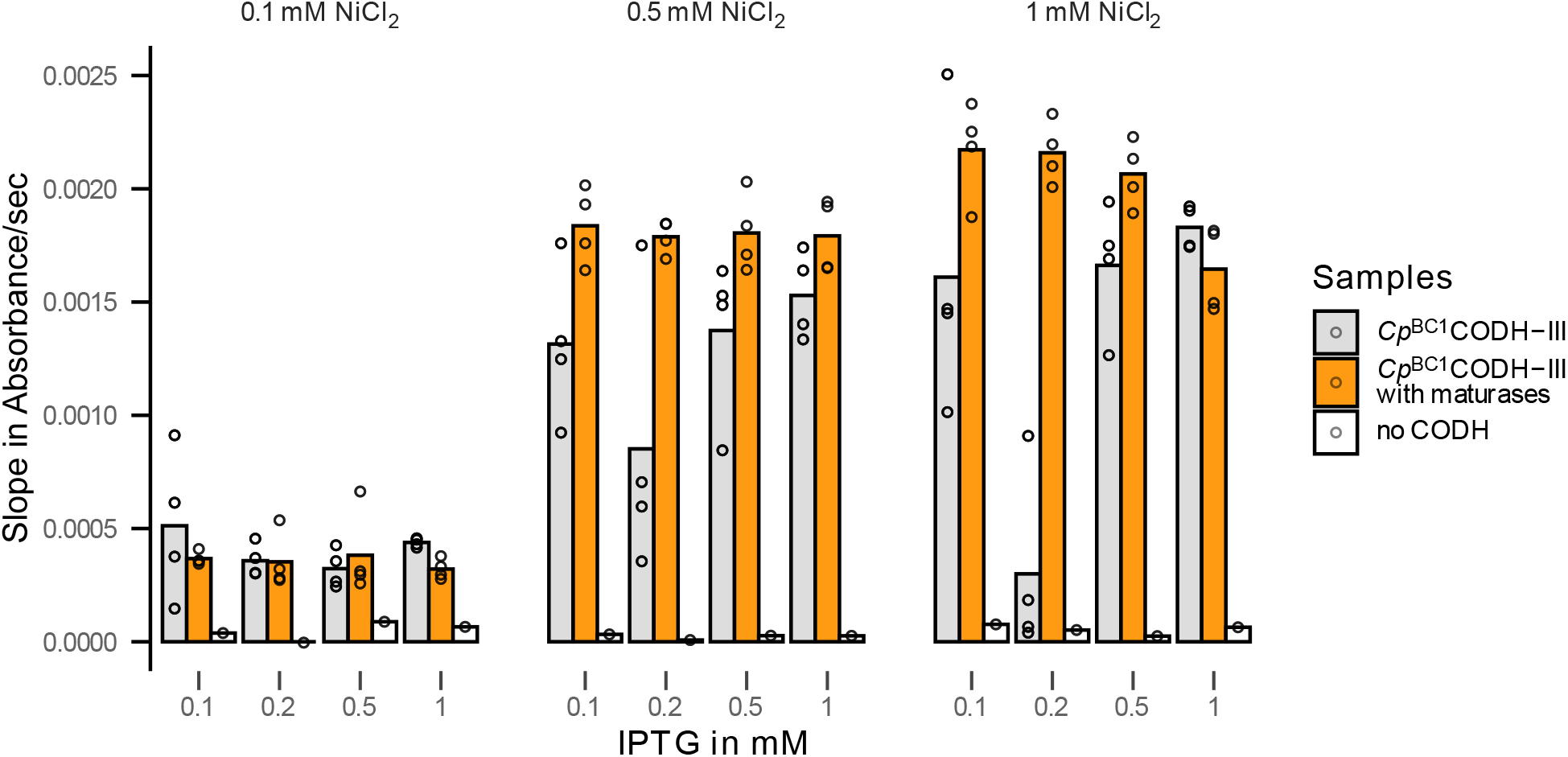
Activity screening of Cp^BC1^CODH-III expressed in combination with maturases (CooCTJ) in orange or without in gray. Each data point is the activity of distinct E. coli colonies. The colonies were grown in a 96-well plate and induced. Activity data was obtained from cleared cell lysate. Lowest and highest value for each condition was removed. Bars represent mean activity.

In order to further study maturase dependence, we compared the operons encoding *Cp*^BC1^CODH-III, *Rr*CODH and CODH from *Shewanelle fodinae* (*Sf*CODH). This comparative analysis aimed to identify conserved features and potential functional divergences among CODHs from clades E and F encoding the CooCTJ operon. *Rr*CODH was chosen as it is the best studied CODH that is encoded together with all three maturases and *Sf*CODH was chosen due to its association with *Sf*CooJ that has been subject to earlier investigation and shows higher similarity to *Cp*^BC1^CooJ.^[6]^ We found that *Rr*CODH and *Rr*CooC have highly similar sequences to *Cp*^BC1^CODH and *Cp*^BC1^CooC-II, and AlphaFold3 (AF3) models predicted similar structures (1.09 RMSD for CODH, AF3 model vs PDB ID:1JQK^[13]^; and 1.54 RMSD for CooC, only AF3 models). The low RMSD values indicate a high degree of structural conservation, suggesting that the core catalytic and maturation functions are preserved across these enzymes. Values describing similarity are summarised in Figure 4. High similarity is also evident by the dot plot in Figure 4, which compares two protein sequences by indicating their similar regions on a matrix like plot. *Rr*CooT/*Cp*^BC1^CooT showed higher variation on a sequence level, but conserved the overall fold as predicted by AF3 (2.01 RMSD, AF3 model vs PDB ID:5N76^[14]^). CooTs nickel binding residue Cys2 is also conserved in *Cp*^BC1^CooT. ^[15]^ Interestingly, a dot plot did not yield matches between the sequences with selected thresholds and window size (Figure 4), supporting the hypothesis by Alfano *et al*. that the N-terminal Cys2 is the most important part of this nickel chaperone.^[15]^ Reduction of both window size and threshold yielded matching pairs, however with no obvious biological meaning (data not shown). This result suggests that while the overall sequence may vary, the functional core—particularly the nickel-binding site—remains conserved.

**Figure 4.**
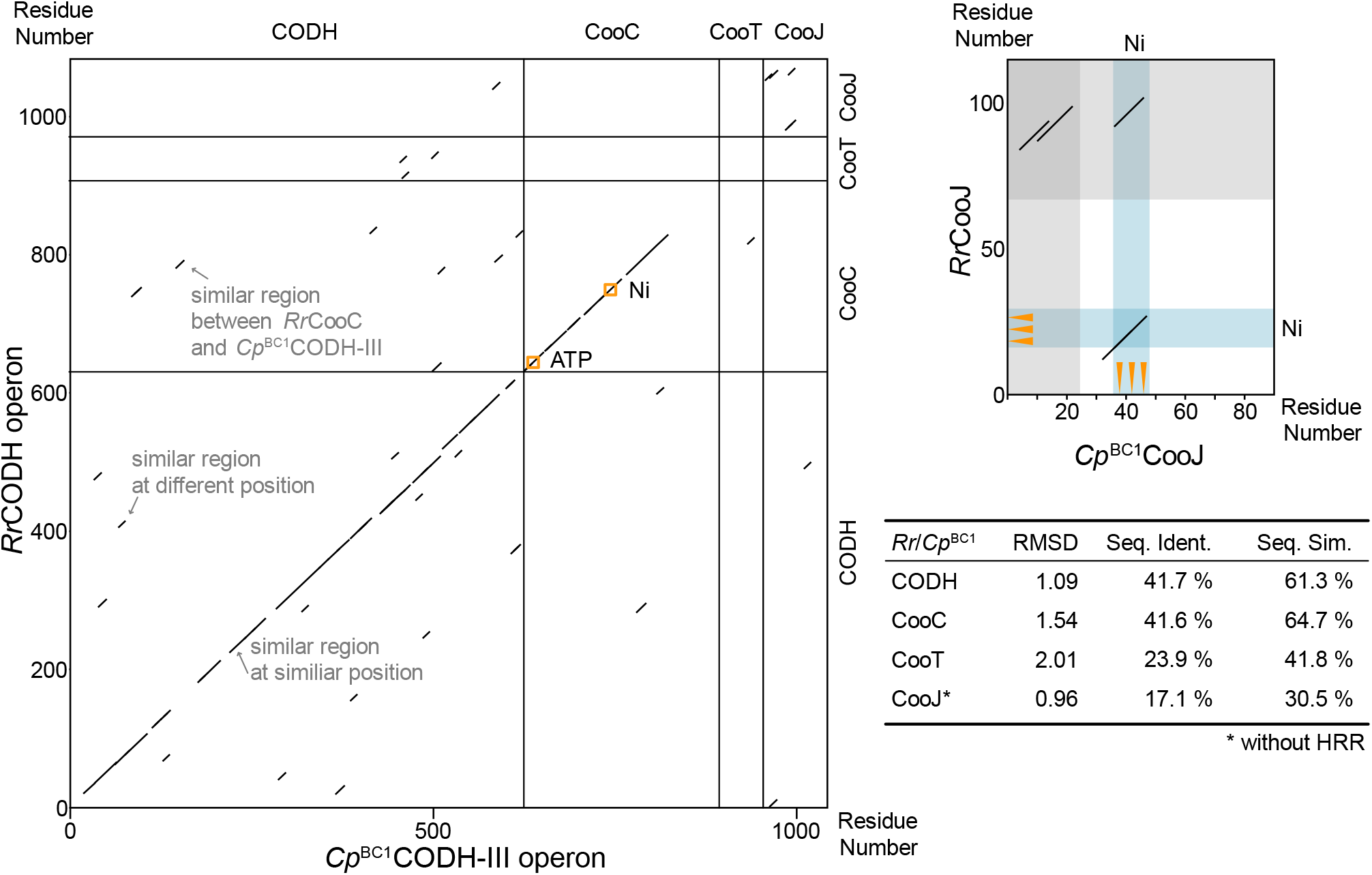
Dot plot of *Cp*^BC1^CODH-III’s operon and *Rr*CODH’s operon. Plots were created with Dotmatcher EMBOSS v6.6.0.0, window-size 10, threshold 23. ATP and nickel binding sites in CooC are marked with orange squares. Gray areas in the CooJ plot are structurally unresolved regions. Blue areas marked with Ni, are metal binding sites in the two proteins. Positions of actual metal binding residues are marked with orange arrows. Table insert summarises the similarity metrics.

Most notably, *Rr*CooJ/*Cp*^BC1^CooJ scored the lowest on all sequence similarity but showed high structural similarity, when aligned without the unordered histidine rich region (HRR) (0.96 RMSD, AF3 model vs PDB ID:6HK5^[7]^). We could identify high similarity regions via the dot plot method (Figure 4), e.g., a metal binding site with the -HWXXHXXXH-motif reported by Darrouzet and co-workers.^[6]^ This conserved region aligns well with the known metal binding sites of *Rr*CooJ, and the AF3 model predicted metal binding sites. Additionally, stretches of similarity could be found in the unstructured HRRs of *Rr*CooJ and *Cp*^BC1^CooJ although at opposite termini. This region could not be modelled via AF3, and was truncated for structural analysis. The inability to model the HRR may reflect its intrinsic disorder, which could be important for its flexibility and interaction with other proteins or metal ions. The presence of similar HRRs, despite their positional differences, suggests a functional convergence in nickel handling, even if the structural context varies.

The same analysis was utilised to compare *Cp*^BC1^CODH/CooCTJ with CODH/CooCTJ from *Shewanella fodinae. Sf*CODH serves as yet another clade F CODH comparison to our clade E *Cp*^BC1^CODH, which also contains the characteristic CODH/CooCTJ operon architecture. *Sf*CooJ was subject to investigation by Darrouzet *et al*. and is more similar to *Cp*^BC1^CooJ, since it also contains its HRR at the N-terminus.^[6]^ For the whole operon we observed similar trends as for *Rr*CODH/CooCTJ with the exception of a higher sequence identity and similarity for *Sf*CooJ. Dot plot analysis showed the conservation of the metal binding site (Figure 5), and another rather conserved stretch in the unordered HRR. CooTs nickel binding residue Cys2 is also conserved in *Sf*CooT.^[15]^

**Figure 5.**
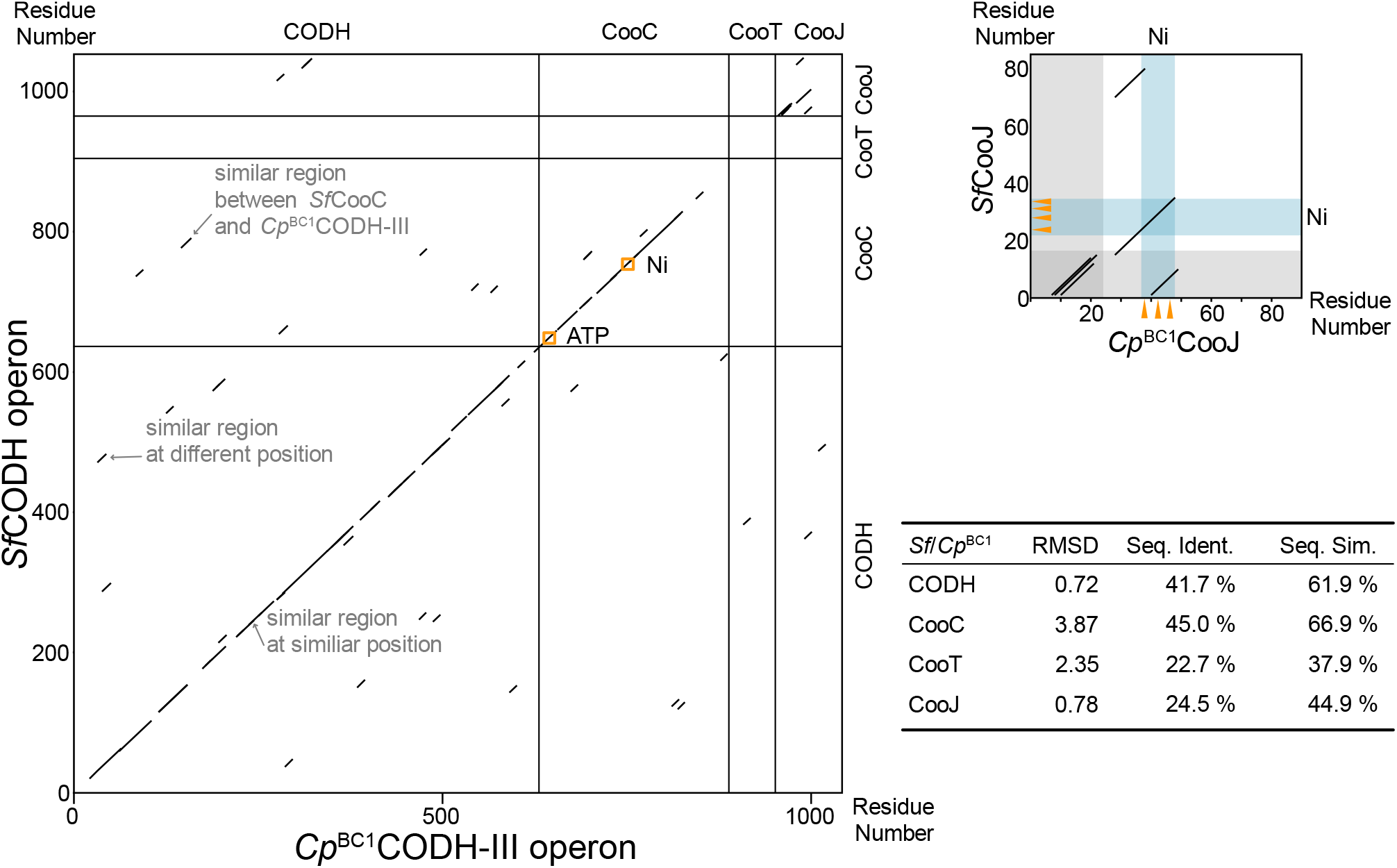
Dot plot of *Cp*^BC1^CODH-III’s operon and *Sf*CODH’s operon. Plots were created with Dotmatcher EMBOSS v6.6.0.0, window-size 10, threshold 23. ATP and nickel binding sites in CooC are marked with orange squares. Gray areas in the CooJ plot are structurally unresolved regions. Blue areas marked with Ni, are metal binding sites in the two proteins. Positions of actual metal binding residues are marked with orange arrows. Table insert summarises the similarity metrics. Note that all RMSD are based on AF3 models since no experimental structural information is currently available.

## Conclusion

The newly characterised CODH-III from *C. pasteurianum* BC1 has similar spectroscopic and catalytic properties compared to other CODH from clade E such as *Nv*CODH.^[11]^ However, its operon structure is more reminiscent of clade F CODHs, like *Rr*CODH suggesting a unique evolutionary trajectory.^[12]^ We could show that if expressed heterologously together with its maturases, *Cp*^BC1^CODH-III is produced in a more stable and reproducible fashion. Yet, unlike other CODHs, its maximum activity is not strictly dependent on the presence of maturases, but is instead more influenced by the amount of nickel available to the protein. This observation supports the hypothesis that this clade E CODH may have recently evolved to acquire maturation machinery primarily to stabilise nickel homeostasis, rather than as an absolute requirement for activity. The maturases likely serve to buffer fluctuations in nickel availability, ensuring consistent activation across diverse environmental conditions.

When comparing the *Rr*CODH operon to the *Cp*^BC1^CODH-III operon, we found that the CODH, CooC and CooT are rather similar, even though CooT was the least similar of the three. CooJ, on the other hand, showed a greater difference between the two operons. Notably, one of the overlapping regions was structurally unresolvable in *R*rCooJ, and is in the C-terminal position, whereas in *Cp*^BC1^Coo ‘s F model it was also not resolved, but is in the N-terminal position, more similar to *Sf*CooJ. The other similar region is the metal binding site characteristic for CooJ with an additional stretch of alpha-helix. CooJ has been reportedly difficult to identify via sequence alone because only small parts of the protein are somewhat conserved.^[4,6]^ While many other CODHs from clade E are encoded with the three maturases CooC, CooT and CooJ, *Cp*^BC1^CODH-III is a unique example that structures them in a CooCTJ fashion.

The discovery of an active Clade E CODH within a Clade F genomic environment expands the known diversity and adaptability of the CODH family. The ability of *Cp*^BC1^CODH-III to assemble and function without its native maturases suggests that the enzyme possesses intrinsic robustness. It has previously been shown that some CODHs do not require their associated maturases;^[16,17]^ however, this is the first example of a CODH containing CooCTJ in its operon that is functionally expressed without them. This finding challenges the traditional view of maturase dependency and opens new avenues for exploring the evolutionary pressures shaping CODH maturation.

Future studies should explore the biochemical consequences of this arrangement, including structural determination, metal binding capacity, and physiological roles in *C. pasteurianum* BC1. *Cp*^BC1^CODH-III thus provides a valuable model to study CODH maturation and evolution, offering valuable insights into the adaptability of these enzymes and their potential applications in biotechnology.

## Material and Methods

Chemicals and enzymes were purchased from Sigma-Aldrich if not stated otherwise.

### Plasmids

*Cp*^BC1^CODH-III’s amino acid se uence (WP_015 15 15.1) and its maturases (WP_015 15 1.1, WP_015615317.1, WP_015615318.1) were codon optimised for production in *E. coli* via GenScript and cloned in pET-11a (restriction sites NdeI and BamHI) and pCDFDuet-1 (restriction sites NcoI and BamHI) vectors respectively. We used a dual vector approach with pET-11a vector carrying *Cp*^CB1^CODH-III and pCDFDuet-1 carrying CooC, CooT and CooJ, within its multiple cloning site 1. The three maturases were separated by a stop codon and a ribosome binding site (5’-AAAGAGGAGAAATTTATA-’). The plasmids were transformed in *E. coli* BL21 DE3 Δ*iscR*.

### Enzyme Purification

For purification the cells were grown to a OD_600_ of 0.5 in phosphate buffered LB medium (20 g/L LB (L3022), 10 mM phosphate buffer, pH 7.4, 1 mM NiCl_2_, 4 mM L-cysteine, 2 mM ammonium iron citrate, 100 mg/L ampicillin, 50 mg/L kanamycin, 50 mg/L streptomycin) at 37 °C while shaking (150 rpm). Then the flask was sealed with a butyl rubber septum, and purged with N_2_ gas for 30 min. During purging, sodium fumarate (final concentration 5 mM) and isopropyl β-D-1-thiogalactopyranoside (IPTG, final concentration 0.2 mM) was added. The cells where then grown over night at 20 °C while shaking (150 rpm). The cells were harvested inside a glovebox via centrifugation (3700 rpm, 20 min), and lysed using lysis buffer (50 mM Tris-HCl, 20 mM NaCl, 2 mM dithiothreitol (M109, VWR Life Science), pH 8, 5 % glycerol, 1 % saccharose, 0.1 % sodium deoxycholate, 1 tablet/50 mL protease inhibitors (05892791001, Roche), 2 mM dithiothreitol, 20 mg/mL lysozyme (62971), 0.5 mg/mL DNAse (DN25), 0.5 mg/mL RNAse (R5503)) and sonication. The lysate was ultracentrifuged for 1 h at 200 000 g. The soluble fraction was syringe filtered and applied to a StrepTrap XT column (Cytiva). The protein was eluted using 50 mM biotin, 50 mM Tris-HCl, pH 8, 20 mM NaCl, after which it was concentrated using 100 kDa cut-off filters, to about 0.5 mM monomer concentration and stored at 4 °C in a glovebox until further use.

### FeS-cluster Reconstitution

Iron-sulphur clusters of the protein were reconstituted semi-enzymatically. The enzyme was diluted in buffer (50 mM Tris-HCl, pH 8, 20 mM NaCl) to a concentration of 50 µM, and dithiothreitol (M109, VWR Life Science), L-cysteine, and ohr’s salt were added in excess (1.5e the amount of iron atoms to be added in the enzyme). The reaction was started by adding 0.01eq cysteine desulphurase (CsdA). The reaction was followed via UV-vis. After no change in absorbance could be observed, the reaction was stopped by buffer exchange using a PD-10 column to plain buffer (50 mM Tris-HCl, pH 8, 20 mM NaCl). The protein was concentrated using 100 kDa cut-off filters to around 0.5 mM monomer concentration and stored at 4 °C in a glovebox until further use.

### Cuvette Assays

The CO oxidation activity of *Cp*^BC1^CODH-III was measured using methyl viologen (MV) as an electron acceptor. In a gas tight cuvette 2.5 mM MV and 10µM sodium dithionite (NaDT) were mixed with buffer (50 mM Tris-HCl, 20 mM NaCl, pH 8) to yield 1 mL reaction mixture. 5µL of enzyme (0.44 µM) was then added to the reaction mixture. The reaction was initiated by adding 25 µL CO saturated buffer and the absorbance was followed at 604 nm.

The CO_2_ reduction activity of *Cp*^BC1^CODH-III was measured using haemoglobin (Hb) as reporter for CO formation, MV as an electron shuttle, and NaDT as sacrificial electron donor. In a tight cuvette 250 µM MV, 2 mM NaDT, 0.1 mg/mL Hb and 10 mM NaHCO_3_ were mixed in buffer (50 mM Tris-HCl, 20 mM NaCl, pH 8) to a final volume of 1 mL. The reaction was initiated by adding 10 µL of enzyme (4 µM), and the absorbance was followed at 429 nm.

### Electron Paramagnetic Resonance spectroscopy

The EPR samples were prepared in an anaerobic glovebox. Protein (100 µM) was incubated with different redox agents for 30 min at room temperature in buffer (50 mM Tris-HCl, 20 mM NaCl, pH 8). Redox agents were: CO saturated buffer, 1 mM NaDT, and 10 mM NaHCO_3_. Alternatively, no redox agents were added to the buffer, this is marked as “as prepared”. fter that, the samples were flash frozen in a -80 °C iso-propanol bath, following a slow transfer to storage in liquid nitrogen. The samples were measured in a Bruker X-Band ELEXYS E500 EPR spectrometer with a SuperX EPR049 microwave bridge and a cylindrical TE011 ER 4122SHQE cavity, coupled to an Oxford Instruments continuous-flow cryostat. All samples were measured at 5 K via liquid-helium flow regulated by an Oxford Instruments ITC 503 temperature controller. Bruker Xepr software package was used to operate the system and export spectra.^[18]^

### Plate Assay

For screening of different expression conditions, *E. coli* BL21 DE3 Δ*iscR* were either transformed with both aforementioned plasmids, only with CODH or with no plasmid. From each transformation, six colonies were selected and grown over night in LB. The next day they were used to inoculate a 96-well plate containing phosphate buffered LB (20 g/L LB (L3022, Sigma-Aldrich), 10 mM phosphate buffer, pH 7.4) with varying concentration of NiCl_2_, 2 mM ferrous ammonium citrate and 4 mM L-cysteine to an OD_600_ of 0.05. The plates were then incubated at 37 °C (750 rpm) for 3 h or until an OD of 0.6 was reached in well A1 of each plate. The plates were transferred to an anoxic atmosphere, OD was measured and expression was induced with varying amounts of IPTG. The plates were than incubated over night at room temperature while shaking (750 rpm). The OD was measured again, before the plates were centrifuged at 3700 rpm for 20 min. Supernatant was removed and lysis buffer (see above) was added. The cells where lysed under shaking (750 rpm) for 1 h. The lysate was then centrifuged (3700 rpm for 20 min) again to separate the insoluble fraction. The soluble fraction was transferred to a new plate. The activity of CO_2_ reduction was assayed. The assay mixture was prepared and stored in a tube before adding 180 µL to 20 µL of crude lysate in a 96 well plate. The assay mix contains: 15 mL buffer (50 mM Tris-HCl, 20 mM NaCl, pH 8), 223 µL MV (25 mM), 445 µL NaDT (100 mM), 2.23 mL haemoglobin (H2500) (2 mg/mL) and 2.23 mL NaHCO_3_ (100 mM) The change in absorbance at 429 nm was followed using a Tecan Infintie 200 Pro MNano plate reader.

### Bioinformatics

For bioinformatic analysis the following sequences were used: *Cp*^BC1^CODH-III (WP_015615315.1); *Cp*^BC1^CooC-II (WP_015615316.1); *Cp*^BC1^CooT (WP_015615317.1); *Cp*^BC1^CooJ (WP_015615318.1); *Rr*CODH (WP_011389181.1); *Rr*CooC (AAC45124.1); *Rr*CooT (AAC45125.1); *Rr*CooJ (AAC45126.1); *Sf*CODH (WP_133040587.1); *Sf*CooC (WP_133040588.1); *Sf*CooT (WP_133040589.1); *Sf*CooJ (WP_133040590.1). All bioinformatic operations were carried out with the EMBOSS software package v6.6.0.0.^[19]^ Dot plots were generated using Dotmatcher with a window-size 10 and threshold 23. For CooT pairing different parameters were screened with no meaningful result (data not shown). Pairwise alignment was done with Needle, using the Needleman-Wunsch global alignment, a gap opening penalty of 10 and a gap extension penalty of 0.5 was chosen.^[20]^ The AlphaFold3 models were generated using the web server.^[21,22]^ For *Sf*CODH and *Cp*^BC1^CODH-III a dimer was modelled. For all CooCs, a dimer with two ATP and one zinc ion was modelled. For *Sf*CooT and *Cp*^BC1^CooT a homodimer was modelled with two zinc ions, and for *Sf*CooJ and *Cp*^BC1^CooJ a truncated version (only residues 29 to 90) with two zinc ions was modelled as a homotetramer.

## Acknowledgements

The Novo Nordisk Foundation (Grant reference number NNF21OC0066716) is gratefully acknowledged for funding.

## Notes

### Competing Interest Statement

The authors have declared no competing interest.

